# Acute effects of prolactin on hypothalamic prolactin receptor expressing neurones in the mouse

**DOI:** 10.1101/2020.05.10.087650

**Authors:** T. Georgescu, S.R. Ladyman, R.S.E. Brown, D.R. Grattan

## Abstract

The anterior pituitary hormone, prolactin, is a fundamental regulator of lactation, and also plays a role in many other physiological processes including maternal behaviour, reproduction, immune response and even energy balance. Indeed, prolactin receptors (Prlr) are widely distributed throughout the body, including a number of different brain regions, further attesting to its pleiotropic nature. Within the brain, previous research has identified key areas upon which prolactin exerts effects on gene transcription through the canonical JAK2/STAT5 pathway downstream of the Prlr. In some neurones, however, such as the tuberoinfundibular dopamine neurones that control prolactin secretion, prolactin can also exert rapid actions to stimulate neuronal activity. While prolactin-induced activation of STAT5 has been described in a wide variety of brain regions, its capacity for acute modulation of electrical properties of many Prlr-expressing neurones remains to be elucidated. To investigate how widespread these rapid actions of prolactin are in various Prlr-expressing neurones, we utilised a transgenic mouse line in which Cre recombinase is specifically expressed in the coding region of the prolactin long form receptor gene (*Prlr-iCre*). This mouse line was crossed with a Cre-dependent calcium indicator (GCaMP6f) transgenic mouse, allowing us to visually monitor the electrical activity of Prlr-expressing neurones in *ex vivo* 200μm brain slice preparations. Here, we survey hypothalamic regions implicated in prolactin’s diverse physiological functions such as: the arcuate (ARC) and paraventricular nuclei of the hypothalamus (PVN), and the medial preoptic area (MPOA). We observe that in both males and virgin and lactating females, bath application of prolactin is able to induce electrical changes in a subset of Prlr-expressing cells in all of these brain regions. The effects we detected ranged from rapid or sustained increases in intracellular calcium to inhibitory effects, indicating a heterogeneous nature of these Prlr-expressing populations. These results enhance our understanding of mechanisms by which prolactin acts on hypothalamic neurones and provide insights into how prolactin might influence neuronal circuits in the mouse brain.

## Introduction

The hormone prolactin is synthesised in the anterior pituitary and has a wide range of physiological and behavioural functions. The regulation of lactation is perhaps prolactin’s best-characterised role, yet its >300 diverse actions include many other processes such as energy balance, stress responses, fertility, and maternal behaviour [1–4]. Consistent with these pleiotropic qualities, the distribution of the prolactin receptor (Prlr) spans numerous peripheral organs and brain regions, including those known to modulate the aforementioned functions [5,6].

The Prlr is a member of the class1 cytokine receptor family, and as such, the primary intracellular signalling cascade responsible for prolactin’s actions is the janus kinase/signal transducer and activator of transcription (JAK/STAT) pathway, with STAT5 being most important for prolactin action [1,4]. Binding of prolactin to its receptor exerts co-ordinated effects on gene transcription dependent on phosphorylation of the STAT5 transcription factor. Utilising immunohistochemical labelling of phosphorylated STAT5 (pSTAT5) has therefore been an effective method of mapping prolactin-responsive cells [7–10]. In the central nervous system (CNS), prolactin-induced pSTAT5 is widely distributed in forebrain and brainstem areas [11]. In some neuronal populations, however, prolactin has also been shown to exert rapid, direct actions affecting electrical activity of the cells [12–19]. The best characterised population are the tuberoinfundibular dopamine (TIDA) neurones in the arcuate nucleus (ARC) of the hypothalamus, which are involved in the feedback regulation of prolactin secretion. In these neurones, prolactin induces a rapid depolarisation and increased rate of firing of action potentials, resulting in increased dopamine release and inhibition of further prolactin release [12,16,20]. In male rats, prolactin drives TIDA neurones from their archetypal oscillations to tonic firing [12,21]. This prolactin-induced depolarisation is postsynaptic and mediated through a transient receptor potential (TRP)-like channel [12]. Similarly, in males and both virgin and lactating female mice, prolactin increases the firing rate of ARC dopamine neurones [16,20]. In these same neurones, transcriptional actions of high prolactin drives changes in gene expression that can influence the neurotransmitter output from the neuron. For example, in males and non-pregnant females, prolactin stimulates expression of tyrosine hydroxylase to promote dopamine synthesis [22]. In late pregnancy and lactation, however, chronic prolactin action promotes enkephalin production from these neurones while dopamine synthesis is reduced [23]. Thus, prolactin action has the potential to both rapidly modify the activity of neuronal circuits, and to induce long-term plasticity in those circuits.

While prolactin-induced activation of STAT5 has been described in a wide variety of brain regions, its capacity for acute modulation of electrical properties of many Prlr-expressing neurones remains to be elucidated. Given that prolactin action is implicated in range of behavioural and physiological adaptations of complex neural circuits, such as expression of maternal behaviour, it is important to better understand how prolactin acts in these different circuits. The aim of the present study was to determine how widespread the rapid actions of prolactin are in different neuronal populations that express the Prlr. Arguably, the most important actions of prolactin in the brain will occur at a time when prolactin levels are elevated, such as during pregnancy and lactation. Hence, we have initially focussed on three prolactin-sensitive hypothalamic regions; the ARC, paraventricular nucleus (PVN), and medial preoptic area (MPOA) [7,8,11], which have been implicated in maternal adaptations during pregnancy, As described above, in the ARC, prolactin is crucial for regulating its own secretion via a short-loop negative feedback acting through TIDA neurones [12,24,25]. Prolactin actions in the ARC also include its effects on fertility, which are modulated via ARC kisspeptin cells [26]. In the PVN, the pleiotropic nature of this hormone is likewise apparent as prolactin has been linked to modulation of oxytocin neurones [15,17,19], the stress response [27,28] and energy balance [29–32]. Some rapid responses to prolactin have been described in the PVN, although responses are heterogenous and differ between virgin and lactating females [15,19]. Perhaps one of the most pivotal maternal adaptation is the ability to successfully care for the offspring. The brain region key for this role is the MPOA, also host to prolactin-activated neurones [7]. Indeed, prolactin elicits marked effects on maternal behaviours through its receptors located in the MPOA [33], with a large number of neurones expressing pSTAT5 in response to prolactin [7,10]. A subset of MPOA neurones are also able to respond electrically to prolactin [18].

We aimed here to enhance our understanding of prolactin’s acute effects by surveying these actions in these three different neuronal populations. To establish cellular specificity, we utilised a transgenic mouse line where GCaMP6f is specifically expressed in Prlr-expressing neurones. By measuring intracellular calcium changes, we examined whether prolactin can induce rapid electrical changes in cells that express the Prlr. As the level of responsiveness in these regions may be altered during periods of sustained elevated prolactin such as lactation [11], we examined both virgin and lactating mice. Since we have recently shown that Prlr expression is also widespread in the male brain [6], we also evaluated acute responses in males.

## Methods

### Animals

The generation and genotyping of Prlr-iCre are detailed elsewhere [6]. Prlr-iCre mice were crossed with Cre-dependent fluorescent calcium indicator GCaMP6f [B6.Cg-Gt(ROSA)26Sor^tm95.1(CAG-GCaMP6f)Hze^] mice from Jackson Labs to generate mice that express GCaMP6f specifically in Prlr-expressing neurones.

All mice were used as adults (8-20 weeks) and were group-housed in a controlled environment: temperature (22 ± 1 °C) and 12h light/dark cycle. All animals had *ad libitum* access to chow pellets and tap water. For virgin female mice, estrous cycle stage was monitored by daily vaginal smears, and animals were used on the morning of diestrous. For lactating groups, the day of parturition was counted as day 1 of lactation, and animals were killed on the morning of day 7 of lactation.

All animal procedures were performed according to the New Zealand Animal Welfare Act and approved by the University of Otago Animal Welfare and Ethics Committee.

### Immunohistochemistry

To assess the expression of green fluorescent protein (GFP) in Prlr-iCre x GCaMP6f animals, male and virgin female mice were terminally anaesthetised with sodium pentobarbital (100 mg/kg, i.p.) and transcardially perfused with 20 ml of 4% paraformaldehyde (fixative). Brains were extracted, post-fixed in a 4% paraformaldehyde fixative solution for 4–6 hours and subsequently transferred to a 30% (wt/vol) sucrose solution in phosphate buffered saline (PBS) for 1-2 days. Brains were sectioned into 30μm coronal sections on a cryostat and collected in 4 equal series. These were stored in cryoprotectant solution at −20°C until the immunohistochemistry was performed.

Free floating sections were incubated for 10 min in 1% H_2_O_2_ (in 0.05 mol L^-1^ Tris-buffered saline, TBS) to block endogenous peroxidases before being blocked for 1 hour in 0.2% bovine serum albumin and 2% normal goat serum. Sections were incubated for 48 hours at 4°C in primary rabbit polyclonal anti-GFP antibody (A-6455; dilution 1: 80 000; Life Technologies, Grand Island, NY, USA). Sections incubated in biotinylated goat anti-rabbit IgG (1:500 BA-1000; Vector Laboratories, Burlingame, CA, USA) for 2 hours at room temperature (RT), before being incubated for 1 hour in avidin–biotin complex (Vectastain Elite ABCKit, Vector Laboratories, Peterborough, UK). GFP immunoreactivity was visualised using glucose oxidase-catalysed nickel-enhanced 3,3′-diaminobenzidine (DAB), resulting in a dark blue/ black precipitate in GFP-positive nuclei and fibres. Sections were mounted onto gelatin-coated slides, air-dried for 90 minutes before being passed through graded alcohols, cleared in xylene and coverslipped with DPX mounting medium.

### Calcium Imaging

Intracellular calcium imaging recordings were made from adult Prlr-iCre x GCaMP6f male and female mice. Mice were injected with a terminal dose of sodium pentobarbital (100 mg/kg, i.p.), decapitated, brains rapidly extracted and submerged in a choline chloride solution containing: 92mM choline chloride, 2.5mM KCl, 1.2mM NaH_2_PO_4_, 30mM NaHCO_3_, 20mM HEPES, 25mM Glucose, 2mM L-Ascorbic acid, 2mM Thiourea, 3mM sodium pyruvate, 10 mM MgCl_2_, 0.5mM CaCl_2_, and bubbled with 5% CO_2_ and 95% O_2_. Coronal brain slices (200μm) were cut using a vibratome (VT1000S; Leica) and incubated for 10 mins at 30°C in the same choline chloride solution containing. Slices were allowed to recover at RT for at least 1 hour in oxygenated HEPES holding solution containing: 92mM NaCl, 2.5mM KCl, 1.2mM NaH_2_PO_4_, 30mM NaHCO_3_, 20mM HEPES, 25mM Glucose, 2mM L-Ascorbic acid, 2mM thiourea, 3mM sodium pyruvate, 1.3mM MgCl_2_, 2.4mM CaCl_2_.

Slices were transferred to a submerged chamber under an upright microscope (Olympus), continuously perfused at a rate of ca. 2 ml/min with oxygenated warm (30°C) aCSF solution containing: 127mM NaCl, 1.9mM KCl, 1.2mM NaH_2_PO_4_, 26mM NaHCO_3_, 10mM Glucose1.3mM MgCl_2_, 2.4mM CaCl_2_. Slices were illuminated through a 40X immersion objective, using the xenon arc light source (300 W; filtered by a GFP filter cube, excitation 470–490 nm; Chroma) and shutter of a λDG-4 (Sutter Instruments). Epifluorescence (495 nm long pass and emission 500 –520 nm) was collected using a Hamamatsu ORCA-ER digital CCD camera. Light source, shutter and camera were controlled and synchronized with the μ-manager 1.4 software. Variations in intracellular calcium concentration were estimated by measuring GCaMP6f fluorescence changes.

A focal plane including at least one fluorescent cell was selected and acquisitions (100-ms light exposure at 2 Hz) were started. Prolactin (250nM, Sigma) was applied in the bath for 5 minutes after a 2 minute control period. At the end of the recordings, the responsiveness of fluorescent cells was tested by bath application of 20mM KCl. For analysis, regions of interest from time series of images were selected in ImageJ. Average fluorescence intensity of in-focus individual somata selections was measured in each frame. A linear regression was used to correct for florescence bleaching using the slope of the signal. Fluorescence intensity data was analysed using Excel and Prism was used to process the data. Relative changes in fluorescence (ΔF/F) were calculated were F is the mean baseline fluorescence intensity for each selected region of interest. Prolactin was considered to have an effect on [Ca^2+^]_i_ if the SD of basal fluorescence was found be smaller than half of the GCaMP6f fluorescence increase. If this criteria was not met, we analysed the effect of prolactin on the number of calcium spikes. For calculating number of calcium events, total spike counts were collected using the pClamp 10 event threshold search.

### Statistics

Statistical analyses were performed using GraphPad Prism 7. All group comparisons were performed using repeated measures multiple comparisons 1way ANOVA (Holm-Sidak posthoc test) unless otherwise stated.

For all experiments, statistical significance was assigned at P < 0.05 and determined using the stated statistical test. Data were presented as mean ± SEM.

## Results

### Prlr-expressing cells are widely distributed through the mouse hypothalamus

In order to visualise and subsequently record the electrical activity of Prlr-expressing cells in our brain regions of interest, we developed the Prlr-iCre x GCaMP6f mouse line by crossing the Prlr-cre mouse line with a GCaMP6f mouse (Figure 1A). We have previously shown that the Prlr-iCre drives expression of Cre-dependent genes specifically in neurones that express the Prlr [6]. To validate the Cre-dependent expression of the GCaMP6f reporter construct, we evaluated the pattern of expression using immunohistochemistry for GFP. Consistent with previous reports, we observed high levels of GCaMP6f expression within the ARC, PVN and MPOA (Figure 1B-D). To investigate the acute electrical effects of prolactin on Prlr-expressing neurones, we bath applied prolactin (250nM) to coronal *ex vivo* ARC, PVN and MPOA brain slices and measured [Ca^2+^] changes as a surrogate of neuronal activity. At the end of each recording, we bath applied KCl (20mM), which served as a way of identifying all of the viable Prlr-expressing cells in a given field of view (Figure 2A-B). A representative example is in Figure 2A-B, showing how fluorescence changes and thereby electrical activity could be measured in all neurones within a particular field of view. In most regions, some cells showed an acute response to prolactin, while others did not show any response, despite being deemed “viable” by showing a response to KCl.

**Figure 1.**
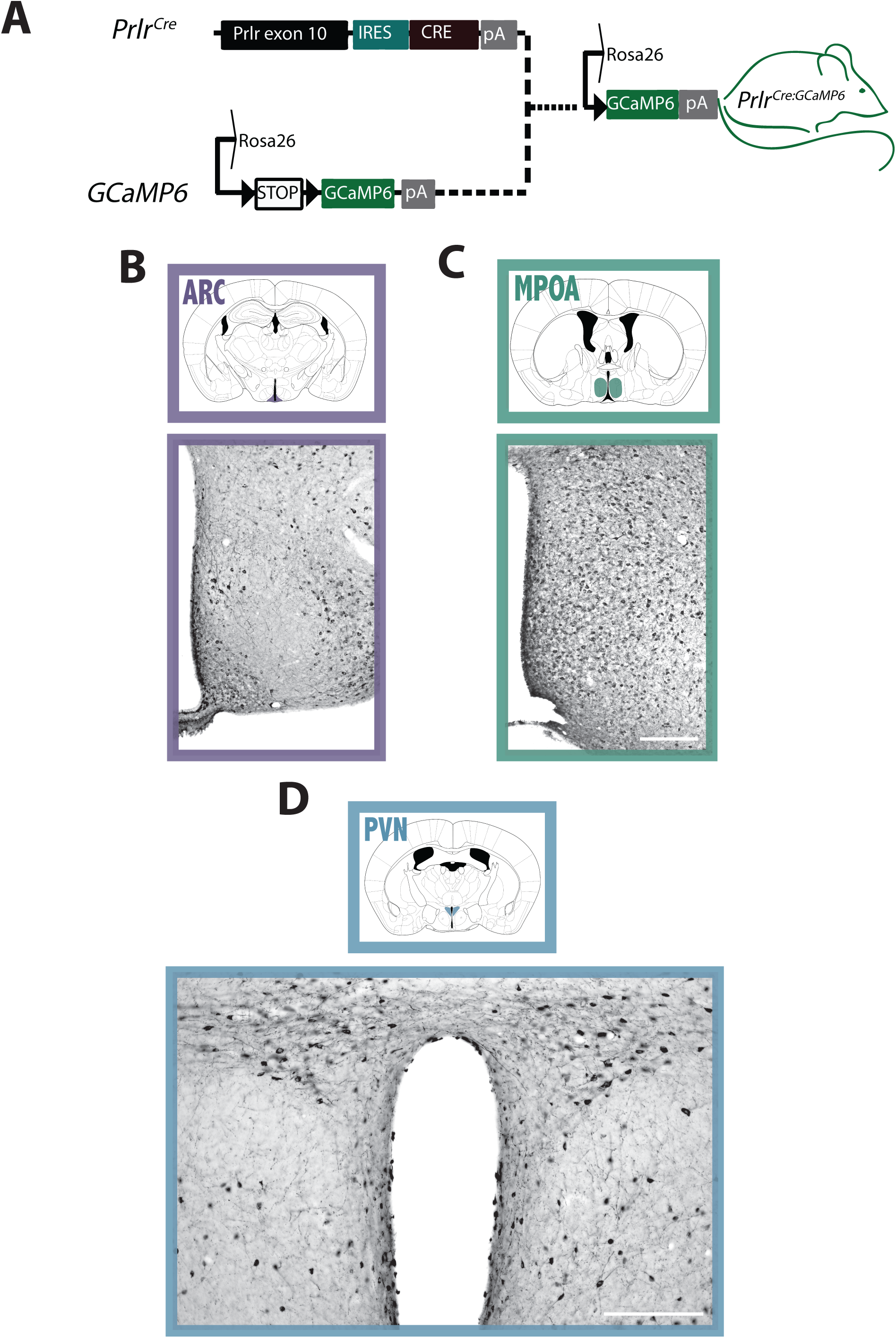
Distribution of Prlr-expressing neurones in the ARC, MPOA and PVN of Prlr-iCre x GCaMP6f mice. A) Schematic detailing generation of Prlr-iCre x GCaMP6f mice. Photomicrographs showing representative Cre-driven GcaMP6f expression (indicated by GFP immunoreactive neurons in black) in coronal brain sections of the ARC (B), MPOA (C) and PVN (D). All diagrams were modified from the Mouse Brain Atlas in Stereotaxic Coordinates, 3^rd^ edition. ARC, arcuate nucleus; MPOA, medial preoptic area; PVN, paraventricular nucleus. Scale bar 200μm.

**Figure 2.**
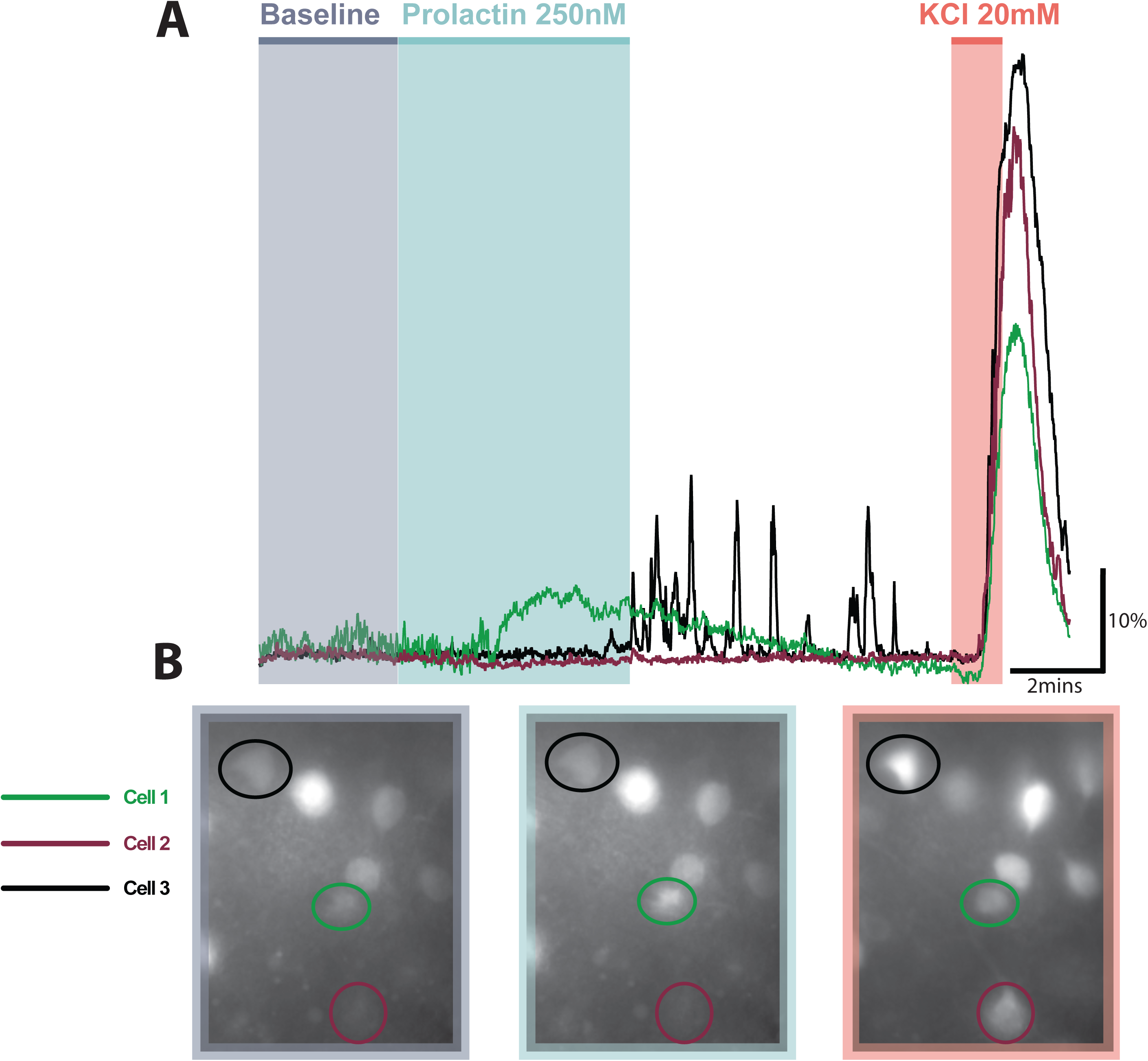
Schemtic illustrating experimental procedure for recording prolactin-induced changes in fluorescence Prlr-iCre x GcaMP6f mice. A) Representative traces from three cells showing different responses to prolactin in an ARC slice from a Prlr-iCre x GcaMP6f mouse. B) Average images (1min bins) during baseline, prolactin application and during the washout period.

### Prolactin activates the majority ARC Prlr-expressing neurones

The ARC is host to a large number of Prlr-expressing cells, widely distributed throughout the extent of this nucleus. Here, we recorded from ARC coronal brain slices of Prlr-iCre x GCaMP6f mice to examine the direct effects of prolactin on these neurones (Figure 3A). In all recordings combined, we observed that prolactin rapidly and reversibly excited a large fraction of ARC Prlr-expressing cells (Figure 3B, 67.36%, 192/285 cells, n=10 animals). In the majority of the cells (62.8%, 179/285 cells, n=10 animals), prolactin induced a significant increase in GCaMP6f fluorescence, which then recovered following the washout period (Figure 3C). A smaller subset of neurones did not show a marked rise in [Ca^2+^], yet we observed a transient increase in the frequency of calcium events (4.56%, 13/285 cells, n=10 animals, Figure 3D-F).

**Figure 3.**
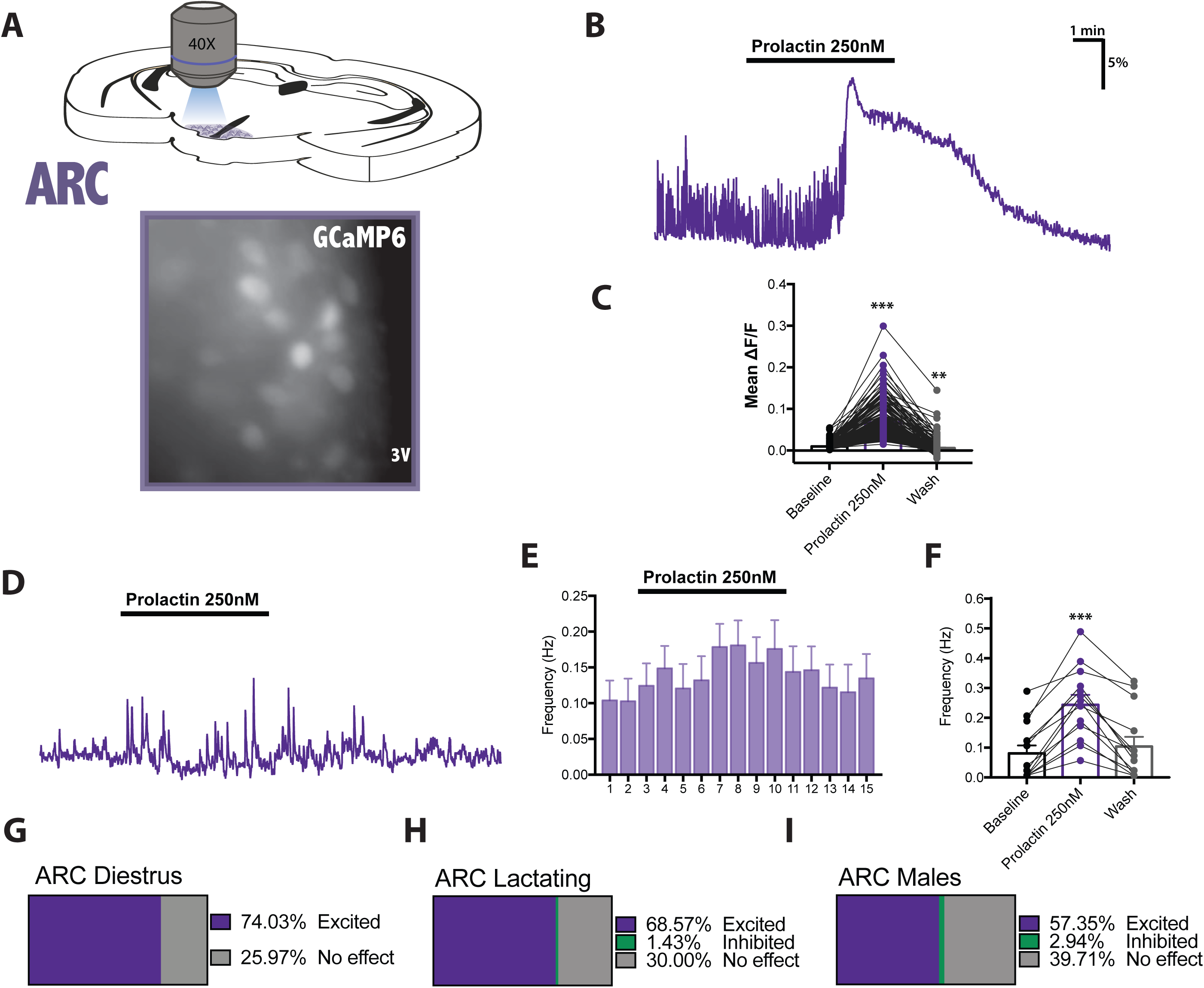
Prolactin activates the majority of ARC Prlr-expressing neurones A) Representative image of a coronal ARC slice from a Prlr-iCre x GCaMP6f mouse. B) Example trace illustrating the excitatory effect of prolactin on the [Ca^2+^] of ARC Prlr-expressing neurones. C) Mean fluorescence changes (1min bins) before, during and after application of prolactin; D) Example trace showing excitatory effect of prolactin on the calcium spike frequency of ARC Prlr-expressing neurones. E) Mean ratemeters (1min bins) of cells showing this excitatory profile; n=13 cells. F) Calcium spike frequency (1min bins) before, during and after the application of prolactin. Proportions of prolactin-responsive ARC Prlr-expressing neurones in virgin diestrous females (G), males (H) and lactating females (I). ***p<0.001 repeated measured 1way ANOVA, vs baseline, Hold-Sidak

The level of prolactin-induced transcriptional activation has been shown to differ under separate physiological conditions or between sexes [11]. For example, lactating dams have elevated pSTAT5 expression in the ARC [11,23]. In addition, Prlrs are expressed in similar numbers in the ARC of males and females [6]. We therefore investigated whether the proportion of prolactin-dependent electrical stimulation differed in diestrous virgin females, lactating females, and males (Figure 3G-I). Our results show that, in all of these three groups, prolactin was able to activate a significant subset of Prlr-expressing neurones: 74.03% of cells were excited in virgin females (Figure 3G, 57/77 cells, n=3 animals), in lactating females 68.57% showed increased activity (Figure 3H, 96/140 cells, n=4 animals), and in males 57.35% of cells were activated (Figure 3I, 39/68 cells, n=3). Conversely, a very small percentage of neurones were inhibited in response to bath application of prolactin: 1.43% in lactating females (2/140 cells, n=4 animals) and 2.94% in males (2/68 cells, n=3 animals), and the remainder of cells were unresponsive.

### A small proportion of PVN Prlr-expressing cells respond to an acute application of prolactin, with a greater level of excitation occurring during lactation

The PVN has a regulatory role in a number of physiological processes that closely overlap with prolactin’s functions [6]. Upon application of prolactin, we observed heterogeneous responses. A subset of PVN Prlr-expressing neurones increased their intracellular [Ca^2+^], thereby becoming more active (18.23%, 45/247 cells, n=15 animals, Figure 4B). The effect of prolactin was mostly reversible as the average fluorescence intensity returned to baseline following the washout (Figure 4C). As in other regions, not all Prlr-expressing cells displayed the same excitation characteristics. We also observed that prolactin caused a transient elevation in calcium event frequency in 10.53% of the cells (26/247 cells, n=15 animals, Figure 4D-F). Prlr-expressing PVN neurones also responded with a reversible inhibition to the acute application of prolactin (8.5%, 21/247 cells, n=15 animal, Figure 4G-I). The level of Prlr expression in the PVN has been shown to be similar in males and females, while lactating females show increased prolactin signalling [6,11]. We therefore sought to establish how the acute effects of prolactin vary between males, virgin diestrous females, and lactating females (day 7 lactation). In diestrous females, 9.62% of cells were excited (5/52 cells, n=4 animals, Figure 4J), 3.85% were inhibited (2/52 cells, n=4 animals, Figure 4J), and 86.54% showed no marked effect (45/52 cells, n=4 animals, Figure 4J). The level of excitation was higher in lactating females (Figure 4K) with 24.09% of cells being activated by prolactin (33/137 cells, n=7 animals) and 15.27% inhibited (12/137 cells, n=7 animals). In males, we found a similar number of cells to be excited as inhibited (7/58 cells, n=4 animals, Figure 4L). Overall, our PVN recordings hint at a heterogenous population that expresses Prlrs, having observed a range of responses.

**Figure 4.**
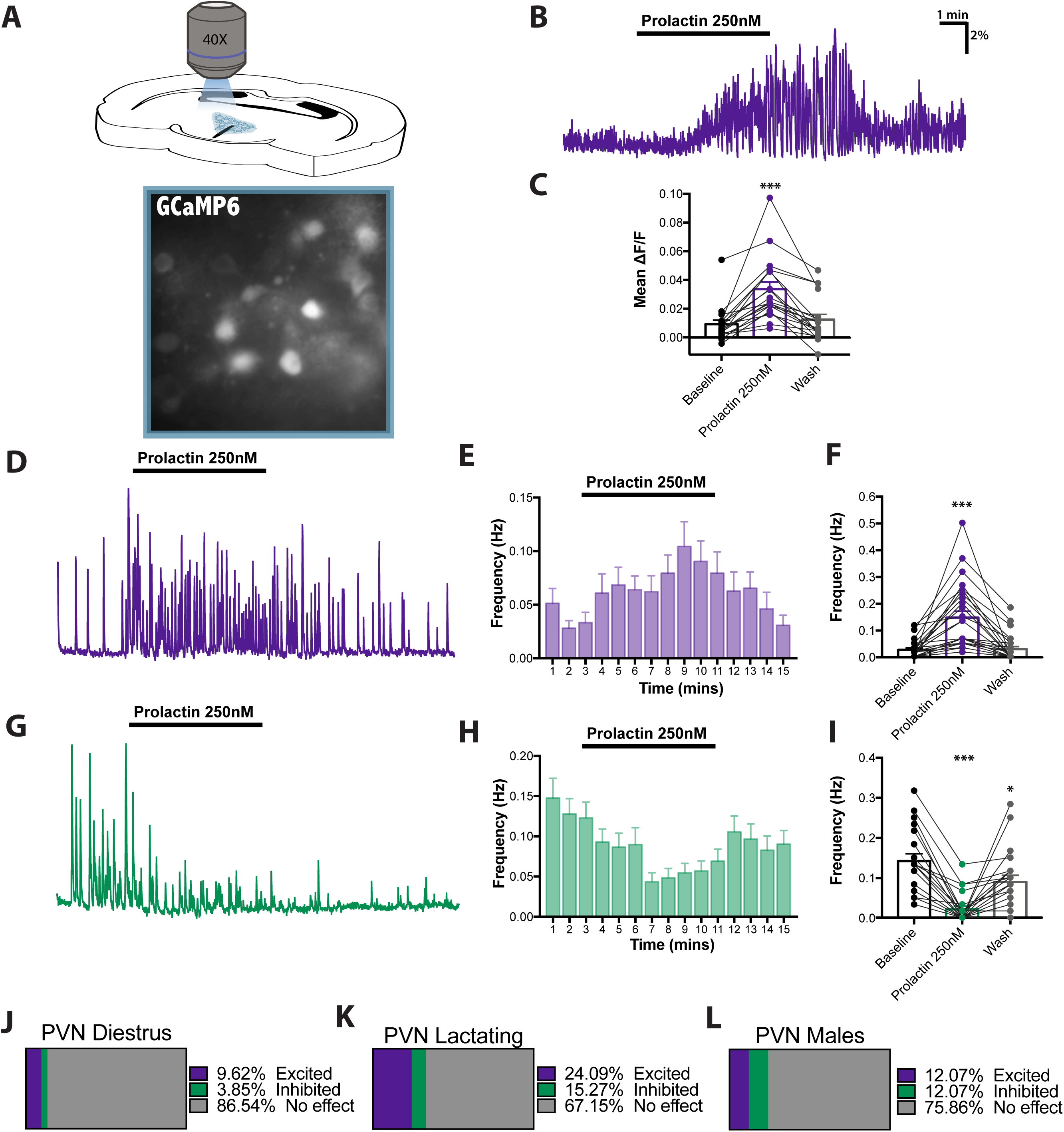
Prolactin has acute effects on a subset of PVN Prlr-expressing neurones A) Representative image of a PVN slice from a Prlr-iCre x GCaMP6f mouse. B) Representative trace showing overall increase in [Ca^2+^] in a PVN Prlr-expressing neurone in response to prolactin. C) Mean fluorescence changes (1min bins) before, during and after application of prolactin. D) Example trace illustrating increase in calcium spike frequency in response to prolactin. E) Mean ratemeters (1min bins) of cells showing this excitatory profile n=26 cells. F) Calcium spike frequency of excited cells (1min bins) before, during and after the application of prolactin. G) Representative trace of prolactin-induced inhibition in PVN Prlr-expressing cells. H) Mean ratemeters (1min bins) of inhibited cells; n=21 cells I) Average calcium spike frequency of inhibited neurones (1min bins) before, during and after the application of prolactin. Proportions of prolactin-induced effects in PVN Prlr-expressing neurones in virgin diestrous females (J), males (K) and lactating females (L). ***p<0.001 vs baseline, repeated measured 1way ANOVA, Hold-Sidak

### Prolactin’s effects on MPOA Prlr-expressing cells

The MPOA is a hub of integration of numerous factors that contribute to parental behaviours, and prolactin action in the MPOA is of particularly relevance for maternal care of the offspring [33]. In response to bath application of prolactin to coronal MPOA slices (Figure 4A), we observed heterogeneous responses. [Ca^2+^] was elevated in a small subset of MPOA Prlr-expressing cells (7.76%, 18/232 cells, n=14 animals) indicating increased neuronal activity (Figure 5B). This depolarisation was reversible as the level of GCaMP6f fluorescence returned to baseline values during the washout period (Figure 5C). To examine whether prolactin can have other rapid effects, we calculated the number of calcium events of cells that were spontaneously active. Bath application of prolactin reversibly increased the frequency of calcium spikes in a small fraction of cells (5.60%, 13/232, n=14 animals, Figure 5D-F). We similarly observed inhibitions in response to the hormone (12.5%, 29/232 cells, n=14 animals, Figure 5G-I). In both of the excited and inhibited groups, prolactin’s effects were transient as the frequency of events returned to baseline recordings following a wash (Figure 4F&I). The proportion of prolactin-induced excitations were similar between virgin females, lactating females, and males (Figure 5 J-L): 13.40% cells in virgin females (13/97 cells, n=5 animals, Figure 5J), 14.47% cells in lactating females (11/76 cells, n=5 animals, Figure 5K) and 11.86% cells in males (7/59 cells, n=4 animals, Figure 5L). Similarly, the level of inhibitions observed did not differ significantly: 13.40% cells in virgin females (13/97 cells, n=5 animals, Figure 5J), 11.84% cells in lactating females (9/76 cells, n=5 animals, Figure 5K) and 11.86% cells in males (7/59 cells, n=4 animals, Figure 5L).

**Figure 5.**
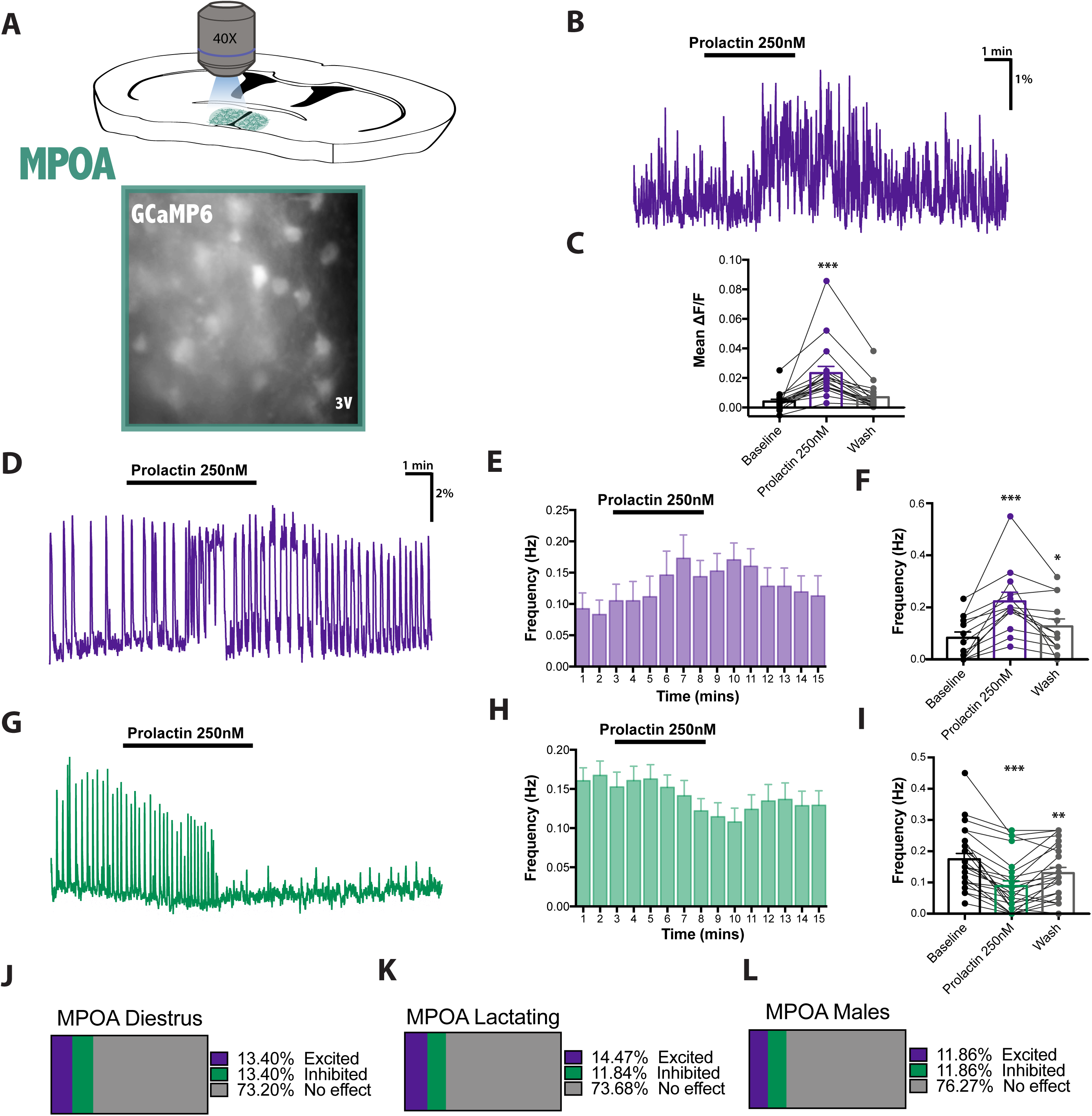
A) Representative MPOA image from a Prlr-iCre x GCaMP6f mouse. B) Representative trace illustrating prolactin-induced increase in [Ca^2+^] in a MPOA Prlr-expressing cell. C) Mean fluorescence changes (1min bins) before, during and after application of prolactin. D) Example trace showing an increase in calcium spike frequency in response to prolactin. E) Mean ratemeters (1min bins) of cells showing this excitatory profile n=13 cells. F) Average calcium spike frequency of excited cells (1min bins) before, during and after the application of prolactin. G) Representative trace of prolactin-induced inhibitions in PVN Prlr-expressing cells. H) Mean ratemeters (1min bins) of inhibited cells; n=21 cells. I) Average calcium spike frequency of inhibited neurones (1min bins) before, during and after the application of prolactin. Proportions of prolactin-induced effects in MPOA Prlr-expressing neurones in virgin diestrous females (J), males (K) and lactating females (L). *** p<0.001 vs baseline; repeated measured 1way ANOVA, Hold-Sidak.

## Discussion

Prolactin is a multifaceted hormone acting through its widely-expressed receptor in the CNS. Identifying the central sites of prolactin action has primarily relied on immunohistochemical labelling for pSTAT5, with very few reports describing rapid electrical effects of prolactin. In the present study, using a genetically encoded calcium indicator as a surrogate measure of neuronal activity, we characterised the acute effects that prolactin has on the electrical activity of Prlr-expressing neurones in the ARC, PVN and MPOA, brain regions that are critical for physiological functions of prolactin. In all these regions, we identified a subset of Prlr-expressing cells that can acutely respond to prolactin. In aggrement with other studies, we found that prolactin rapidly increases the intracellular [Ca^2+^] in the majority of ARC Prlr-expressing neurones, thereby activating them [12,16]. IWe report here that in contrast to the ARC, there are smaller proportions of Prlr-expressing cells in the PVN and MPOA that electrically responded to prolactin. In these brain regions, a similar proportion of inhibitory to excitatory responses occured, compared to the more robust and consistent activation seen in the ARC.

In the ARC, neuronal populations in which prolactin function has been well investigated include the TIDA and kisspeptin neurones. Prolactin secretion is tightly regulated by TIDA neurones that release dopamine at the median eminence to inhibit the constitutively active prolactin release [34–36]. In both rats and mice, upon application of prolactin, TIDA neurones become depolarised and switch from their hallmark oscillation to tonic action potential discharge [12,16,20]. Immunohistochemical methods have revealed a great degree of co-localisation between TIDA neurones and Prlr-expressing cells [6], suggesting prolactin can directly stimulate ARC Prlr-expressing neurones. It seems likely that the majority of the neurones acutely activated by prolactin in the arcuate nucleus will be the TIDA neurones. Consistent with other studies carried out on TIDA neurones [20], we observed little difference in the proportion of prolactin-responsive Prlr-expressing cells between males, virgin females and lactating female mice.

Kisspeptin neurones also reside in the ARC and their activity and function is influenced by prolactin [26,37,38]. Prolactin increases the expression of pSTAT5 in both ARC and AVPV kisspeptin neurones [16,26,37,38], but only has acute effects on 5% of the AVPV population [16]. In the ARC of female mice however, a high majority of kisspeptin cells were found to be prolactin-responsive, as measured through pSTAT5 immunohistochemistry [26]. Specifically, prolactin treatment decreases ARC kisspeptin mRNA expression, raising the possibility that prolactin would inhibit ARC Prlr-expressing neurones [38]. However, we did not see many cells acutely inihibted by prolactin in the ARC, suggesting that this inhibitory action may be predominantly transcriptionally mediated. Alternatively, as the ARC is also host to a cluster of prolactin-resposive GABA-ergic neurones [39], ARC kisspeptin neurones might be indirectly inhibited through acute activation of GABA release. The heterogeneity of the ARC Prlr-expressing population was further underlined by our observation of two categories of excited cells: those that responded with an overall increase in intracellular [Ca^2+^] and those that showed an alteration in calcium spike frequency. Identifying the precise neurochemical identity of these prolactin-responsive neurones would therefore be a interesting avenue for future studies.

The mechanism behind the prolactin-induced acute effects has been explored in only a handful of studies thus far. In sensory neurones in female rats, prolactin increases the activity of TRPV1, TRPA1 and TRPM8 channels [40–42]. These effects are sex-dependent, with prolactin appearing to only regulate nociception in female and not male rodents [42–44]. In male rats, prolactin-induced TIDA neurone activation is post-synaptic as it endures in the presence of the voltage-gated Na^+^ channel antagonist, tetrodotoxin [12]. The response is characterised by a mixed-cationic current, mediated via transient receptor potential (TRP) channels, possibly one of the TRPC channels [12,45]. The high degree of expression of Prlrs in TIDA neurones alludes to the involvement of a similar ion channel in the responses we observed, yet the precise mechanistic underpinnings of these remain to be defined.

Another hypothalamic region where there is convergence of many of the physiological roles of prolactin is the PVN. This key autonomic control centre hosts Prlr-expressing cells which appear to be present in both the parvocellular and magnocellular delimitations of this nucleus [6]. Related to its function in lactation and milk-ejection, prolactin has been shown to interact with PVN oxytocin neurones [17]. Specifically, prolactin hyperpolarises and reduces the firing rate of PVN oxytocin cells in virgin and pregnant but not lactating rats [15,17,19]. In contrast to there reports, our results indicate a different proportion of prolactin-driven inhibitions in Prlr-expressing cells in the mouse. For example, we detected very few cells that were inhibited in virgin female mice while the percentage of inactivations was higher in lactating mice (3.85% in virgin females vs 15.27% in lactating females). These discrepancies may arise due to species differences since the high degree of Prlr expression in oxytocin neurones is not as evident in mice as it is in rats [6,17]. Secondly, our method for detecting inhibitions was reliant on observing a decrease in the frequency of spontaneous calcium events. However, cells lacking spontaneous activity may have not been included in our analysis, resulting in a smaller percentage of inhibitions observed in virgin females. When comparing virgin females to lactating females, previous studies have reported a marked increase in the expression of pSTAT5 in the PVN [11]. Similarly, we found a greater proportion of neurones to be acutely prolactin-responsive in lactating females than in virgin females. Previously unreported, we also detected that approximately a quarter of PVN Prlr-expressing neurones responded in males, with half of those being inhibitions and half excitations. Despite the difference observed between virgin females, lactating females and males, we consistently observed that a vast majority of neurones did not respond to an acute application of prolactin. In contrast to the ARC, this suggests that prolactin induces slower transcriptional responses in more than half of PVN Prlr-expressing cells.

Not dissimilar to the PVN, Prlr-expressing cells of the MPOA were also less responsive to an acute application of prolactin. The MPOA consists of a highly heterogeneic population of neuronal cell types which are closely linked to the regulation of maternal and paternal behaviours [33,46–50]. Indeed, prolactin’s role in offspring care has been attributed to the expression of Prlrs in this brain region [10,33]. Very few studies, however, have examined whether these functions can be attributed to rapid electrical effects or prolactin’s transcriptional signalling. Evidence exists that in the absence of neuronal STAT5, prolactin is able to depolarise approximately a quarter of unidentified MPOA neurones [18]. Our results support earlier stuides that prolactin can induce rapid electrical effects specifically on MPOA Prlr-expressing cells. However, we detected a smaller percentage of cells acutely excited by prolactin than previously reported [18]. The vast neuronal diversity of the MPOA suggests that Prlr-expressing cells however represent only a fraction of neurones present in this brain region. This raises the possibility that prolactin can also depolarise MPOA non-Prlr-expressing neurones via a pre-synaptic mechanism. Similar to the ARC, we noted two subclasses of MPOA excitations, alluding to different types of responsive cells. Contrary to other reports [18], only a small subset of neurones were inhibited in response to bath application of prolactin.

Expression of the Prlr and of pSTAT5 in the MPOA has been reported to be sex-dependent [6,9]. Nonetheless, our investigations detected little difference in the proportion of prolactin-responsive cells in virgin males, females or lactating females. Interestingly, the lack of significant variation also extended to the percentage of inhibitions we observed. This raises the possibility that Prlr-expressing cells that can acutely respond to prolactin subserve similar functions in both virgin males and females and lactating females. Whether this subclass of neurones is characterised by any molecular similarities remains to be determined. In all three groups, the large majority of cells were unresponsive to acute prolactin stimulation, raising the possibility that the transcriptional pSTAT5-dependent pathway is the primary mechanism mediating the actions of prolactin in the MPOA.

Overall, the results presented here have characterised the previously unexplored acute effects of prolactin on Prlr-expressing cells in a number of different hypothalamic regions. We highlight a potentially novel component of the prolactin-signalling pathway by identifying small subsets of neurones that can respond electrically to the hormone in all regions examined. These experiments therefore provide us with an improved appreciation of the complexity of functions engaged by prolactin.

## Acknowledgements

We thank Pene Knowles for genotyping expertise and other technical assistance. We also thank Dr Joon Kim and Dr Richard Piet for assistance with calcium imaging analysis.

